# Diurnal Variations in the Cholinergic System of the Nucleus Accumbens

**DOI:** 10.64898/2026.10.17.759680

**Authors:** Taylor A. Stowe, Yanhua H. Huang, Mark J. Ferris, Colleen A. McClung

## Abstract

The nucleus accumbens (NAc) plays a critical role in reward-related behaviors, which exhibit diurnal variations. Cholinergic interneurons (CINs) are the primary source of acetylcholine within the NAc and regulate local dopamine (DA) signaling through nicotinic acetylcholine receptors (nAChRs). Although there are diurnal variations in NAc dynamics, little is known about whether CIN activity and nicotinic signaling also vary across the light/dark cycle. Here, we aimed to determine diurnal variations in the NAc cholinergic system using *ex vivo* fast-scan cyclic voltammetry and cell-attached electrophysiological recordings in male rodents. We examined time-of-day-dependent differences in nAChR-mediated modulation of DA release, spontaneous CIN firing frequency, and firing patterns. We found that spontaneous CIN firing varied across the light/dark cycle, with increased activity during the dark cycle. Additionally, bath application of nicotine (NIC), mecamylamine (MEC), and dihydro-β-erythroidine (DHβE) reduced CIN firing frequency during the dark cycle, with only MEC decreasing firing frequency in the light cycle. NIC and MEC also altered CIN firing patterns during the dark cycle, whereas DHβE primarily affected firing frequency. Together, these findings demonstrate diurnal variations in NAc CIN activity and reveal time-of-day-dependent differences in the effects of nAChR modulation on spontaneous firing. These results suggest that nicotinic signaling may contribute to diurnal regulation of NAc cholinergic activity, providing insight into mechanisms underlying time-of-day-dependent changes in reward circuitry.

## Introduction

Cholinergic interneurons (CINs) play a role in microcircuits within the striatum, including those within the nucleus accumbens (NAc)^1,2^. Although CINs only make up about 1-2 % of the NAc ^3,4^, they are a primary source of acetylcholine (ACh), which makes them potent modulators of dopamine (DA) signaling ^5^. For example, the removal of endogenous ACh leads to a robust decrease in DA since ACh acts on nicotinic acetylcholine receptors (nAChRs) located on DA terminals to regulate DA release^6^. These receptors are composed of different combinations of α and β subunits, which confer distinct pharmacological and functional properties^7^. β2-containing nAChRs are expressed on DA terminals and contribute to the regulation of DA release, while nicotinic signaling can also influence CIN activity through direct and local circuit-mediated mechanisms. In turn, DA released from these nAChRs acts on inhibitory D2 receptors located on CINs. There is an overall potential bidirectional relationship between ACh/DA, which is a key part of the CIN system that may be mediated by both CIN activity and nAChRs. However, whether these interactions and nicotinic modulation of CIN activity vary across the light/dark cycle remains unclear

CINs are tonically active neurons that exhibit characteristic firing patterns consisting of periods of tonic activity, bursts, and pauses in both *in vivo* and *ex vivo* preparation^8,9^. These patterns can be placed into the following categories: irregular, regular, mixed-mode and burst firing^9^. Tonically active CINs with varying firing patterns are both unique and important for the modulation of NAc dynamics and reward-related behaviors. For example, interfering with ACh release from CINs impaires reward-learning during a Pavlovian task ^10,11^. Better understanding of the CIN system in the NAc may allow us to better parse out the neural mechanisms underlying reward-related behaviors. Specifically, we aim to better understand diurnal (light/dark) differences within the CIN system in the NAc.

The daily, constant solar cycle allows organisms to synchronize circadian rhythms and sleep/wake cycles and lead to diurnal differences in physiology and behaviors^12,13^. For example, body temperature and hormone release fluctuate across the light/dark cycle. Reward-related behaviors also exhibit diurnal variation, including responses to natural rewards in male rats^14^. Moreover, diurnal variations also extend to the neurobiology underlying reward, with light/dark differences observed in both medium spiny neuron (MSN) excitability and dopamine (DA) dynamics, including stimulated DA release, within the NAc^10,15,16^. These neurobiological variations may contribute to time-of-day differences observed in drug taking and craving ^17–19^. Although diurnal variations have been established in local NAc dynamics, our understanding of diurnal variations in the CIN system in the NAc remains limited.

Diurnal variations exist in ACh modulation of DA release^10^; however, whether these differences extend to nAChR modulation of DA release and the intrinsic activity of CINs remains unclear. Here, we aimed to determine diurnal variations across multiple components of the NAc CIN system and investigate potential mechanisms contributing to these differences. We utilized both *ex vivo* voltammetry and electrophysiology to parse out different aspects of the CIN system. First, we determined the diurnal variations in nAChR modulation of DA release in the NAc via *ex vivo* voltammetry. Next, we determined the diurnal variations in tonic CIN activity and firing patterns in the NAc via *ex vivo* electrophysiology. Lastly, we focused on the potential role of nAChRs in the diurnal variations in tonic CIN firing by using pharmacological tools during electrophysiology recordings. These data collectively enhance our understanding of diurnal variations in the CIN system that may play a role in NAc dynamics and reward-related behaviors.

## Methods

### Slice preparation

For voltammetry, male Sprague-Dawley rats were anesthetized with isoflurane and then euthanized by rapid decapitation in a ventilated area free of blood or tissue from previous animals at zeitgeber time ZT 6 (6 hours into light cycle) or ZT 18 (6 hours into dark cycle) depending on group assignment. Rats were always kept in their respective light or dark cycle until brain extraction was complete. These times were chosen based on our previous work determining diurnal variations in stimulated DA release. A vibrating tissue slicer (VF-300 Microtome; by Precisionary Instruments Inc., San Jose, CA) was used to prepare 400 μm thick coronal brain sections containing the NAc core, as previously described (Siciliano et al., 2017). The tissue was immersed in oxygenated, pH-adjusted (7.4) artificial cerebrospinal fluid (aCSF) containing (in mM): NaCl (126), KCl (2.5), NaH2PO4 (1.2), CaCl2 (2.4), MgCl2 (1.2), NaHCO3 (25), glucose (11), and l-ascorbic acid (0.4). Brain slices were transferred to wells containing oxygenated aCSF (32°C) flowing at 1 mL/min.

For electrophysiology, slice preparation varied slightly. Male ChAT-gfp mice were used to specifically target CINs in the NAc and were anesthetized with isoflurane and then euthanized by rapid decapitation in a ventilated area free of blood or tissue from previous animals at ZT 2 (2 hours into light cycle), ZT 6 (6 hours into light cycle), ZT 14 (2 hours into light cycle) or ZT 18 (6 hours into dark cycle) depending on group assignment. Brains were placed into ice-cold oxygenated (95% O2/5% CO2) modified artificial CSF (aCSF) containing the following (in mM): 135 N-methyl-D-glucamine, 1 KCl, 1.2 KH2PO4, 1.5 MgCl2, 0.5 CaCl2, 70 choline bicarbonate, and 10 D-glucose, pH 7.4 adjusted with HCl. NAc-containing coronal slices (250 μm) were sectioned with a vibratome (VT1200S; Leica) and incubated for 30 min at 37°C in oxygenated aCSF containing the following (in mM): 119 NaCl, 26 NaHCO3, 2.5 KCl, 1 NaH2PO4, 2.5 CaCl2, 1.3 MgCl2, and 11 D-glucose. Slices were kept at room temperature until recording then perfused with aCSF (30 –32°C).

### Ex vivo voltammetry

*Ex vivo* fast scan cyclic voltammetry (FSCV) provides unique advantages for studying local circuit modulation of DA. FSCV has high spatial and temporal resolution and allows the examination DA while modeling of a wide range of neuronal firing patterns^20^. Here, ex vivo FSCV was used to determine diurnal rhythms in stimulated DA release in the NAc core across increasing stimulation parameters^21^. A carbon-fiber recording microelectrode (100-200 μM length, 7 μM diameter) was placed 100-150 μM from a bipolar stimulating electrode in the NAc core. DA release was evoked by a single electrical pulse (750 μA, 2 msec, monophasic) applied to the tissue every 3 minutes for a baseline then followed by increasing firing frequencies to mimic phasic-like firing. Extracellular DA was recorded by applying a triangular waveform (−0.4 to +1.2 to −0.4 V vs Ag/Agcl, 400 Vs).

### Ex vivo electrophysiology

*Ex vivo* electrophysiology compliments our previous voltammetry work as we can directly look at a specific cell-type. Male ChAT-gfp mice were used to specifically target CINs in the NAc. Recordings were conducted during the light phase. Slices were viewed by differential interference contrast optics (Leica) and accumbal regions were localized under low magnification. Recordings were made from NAc core and shell CINs under visual guidance with a 40 objective. GFP cells were visualized using filters for 488 nm light, respectively. Borosilicate glass pipettes (3–5 M) were filled with the ACSF for cell-attached recordings. A seal of 15 MΩ or higher was made and only recordings that were stable and had good access to the cell were utilized.

### Bath application of drugs during recordings

Both voltammetry and electrophysiology experiments utilize drugs as pharmacological tools to target nAChRs. For voltammetry, nicotine (NIC; 500 nM), mecamylamine (MEC; 2 µM), or dihydro-β-erythroidine (DHβE; 500 nM) combined with conotoxin (CON) were bath-applied through a perfusion system as previously described in Fennell et al., 2020. DA release was measured following single-pulse and five-pulse electrical stimulation at varying frequencies to determine the effects of nAChR modulation on tonic- and phasic-like DA signaling. Drug-induced changes in stimulated DA release were expressed as a percentage of baseline release and compared between recordings obtained during the light and dark cycles. In our electrophysiology recordings, NIC (1µM), MEC (1µM), and DHβE (10 μM) were bath applied through aCSF perfusion after stable baseline recordings were taken. These doses were chosen based on previous research. Recordings were taken 15 minutes of drug application began.

### Data acquisition and modeling

Demon Voltammetry and Analysis software was used to acquire and model FSCV data ^22^.Recording electrodes were calibrated by recording electrical current responses (in nA) to a known concentration of dopamine (3 mM) using a flow-injection system. This was used to convert electrical current to dopamine concentration ^20^. Electrophysiology recordings lasted for 4 hours and were collected with a MultiClamp 700A amplifier (Molecular Devices) then were filtered at 2.6–3 kHz and amplified 10 times, then digitized at 20 kHz with a Digidata 1322A analog-to-digital converter (Molecular Devices). Both pClamp10 software (Molecular Devices) and DataView (Dr. William Heitler) were used to analyze the spontaneous firing of CINs with a threshold search.

### Statistical analysis

Statistical analyses were performed using GraphPad Prism. For tonic CIN firing, unpaired t-tests were performed. For drug wash on experiments, firing frequency was analyzed using a two-way repeated-measures ANOVA with light–dark phase as the between-subject factor and recording condition (baseline versus drug) as the within-subject factor. The phase × condition interaction was used to determine whether the drug-associated change in firing differed between phases. Baseline and drug firing within each phase were compared using Šídák-adjusted multiple comparisons. Each paired recording was obtained from an independent mouse. Data are presented as mean ± SEM with individual observations shown. Sample sizes and the numbers of animals and cells are provided in the figure legends. Statistical significance was defined as p < 0.05.

## Results

### Diurnal differences in nAChR modulation of stimulated DA release in the NAc

We see in both the light and dark cycle that DA release increases as a % of base across varying frequencies (Fig 1). Our previous found diurnal variations both stimulated DA release and phasic/tonic ratios with higher/phasic tonic ratios in the dark cycle^10^. To expand from these data, we used pharmacological tools to target nAChRs and determine if diurnal variations exist in nAChR modulation of DA release as previously described in both Fenell et al., 2020 and Stowe et al., 2022 (Fig. 1). Nicotine (NIC), a nAChR agonist, desensitizes the receptors at higher doses. This desensitization of nAChRs causes an initial decrease of DA release, but causes higher release at higher stimulating frequencies, leading to an increase in phasic/tonic DA release ratios. Here, we see significant decreases in DA release across stimulation parameters in the light cycle (Fig. 1A) while in the dark cycle, DA release returns to baseline at higher frequencies (Fig. 1B). We followed up on these data using mecamylamine (MEC), a nAChR antagonist, and found a significant decrease in DA release in the light cycle (Fig. 1C) while we see slight decreases at lower frequencies in the dark with an increase at phasic-like firing (Fig. 1D). Lastly, we utilized DHβE, a nAChR antagonist targeting β2-containing nAChRs, in combination with conotoxin (CON) With DhβE + CON, administration we see little to no changes in DA release across firing frequencies in both light (Fig. 1E) and dark cycle (Fig. 1F). Overall, there is a slight increase or return to baseline DA release in the dark cycle after NIC and MEC while DA release remains decreased in the light cycle. However, there may be other local modulators playing a role in mediating diurnal variations in DA release in the NAc.

**Figure 1.**
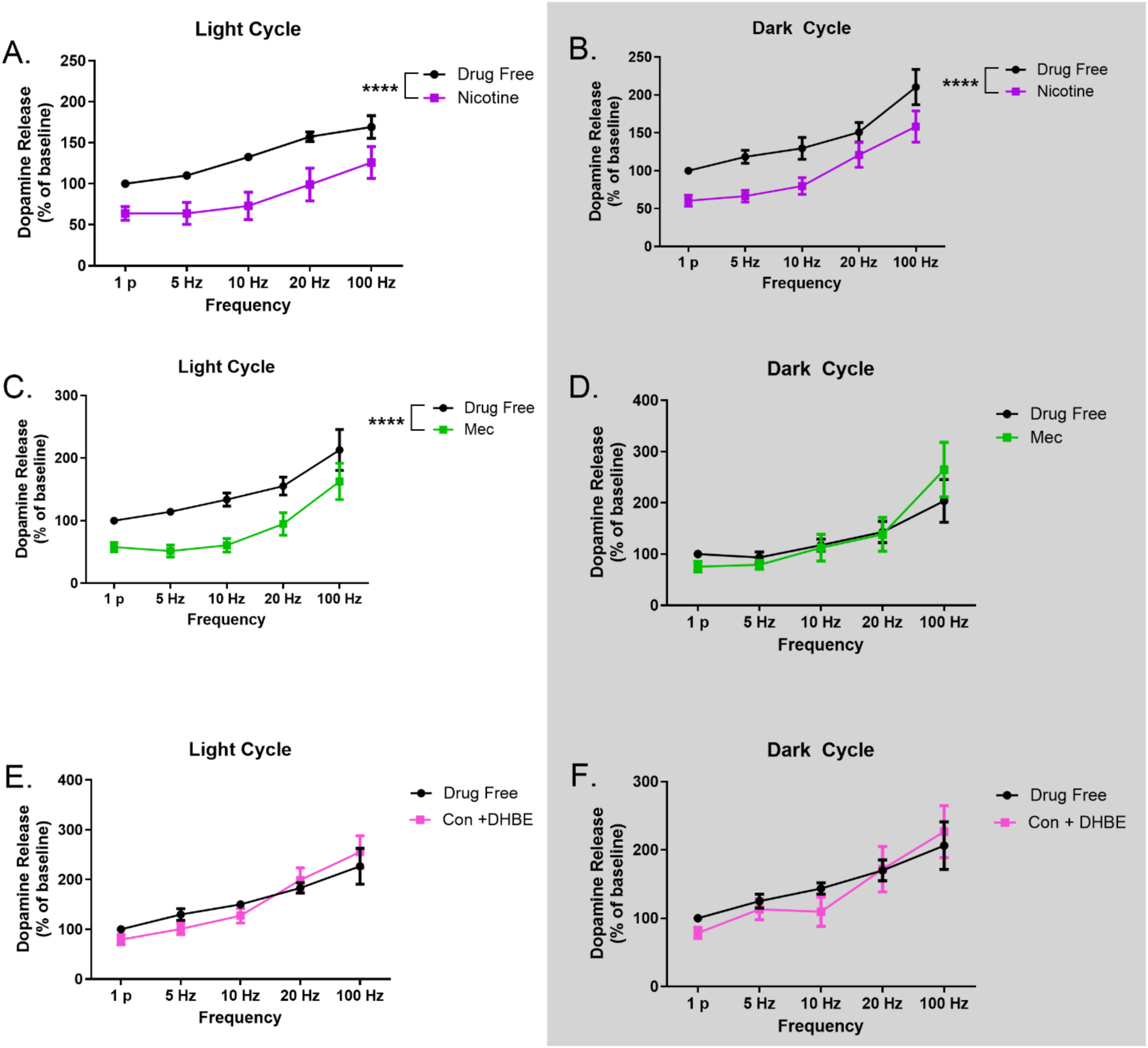
Diurnal variations exist in stimulated DA release across varying frequencies when targeting non-specific nAChR subunits. (A) NIC application decreases DA release in the light cycle across all stimulation parameters (B) In contrast, there is a return to baseline after NIC application in phasic-like frequencies in the dark cycle (C) MEC application decrease DA release across all stimulation parameters (D) MEC has little effect on DA release at lower frequencies in the dark cycle while at phasic-like firing we see a slight increase (E) CON and DHBE had little to no effects on DA release in the light cycle with a slight increase at higher frequencies (F) CON and DHBE, like the light cycle, had little to no effects on DA release as a percentage of baseline with slight increases at higher frequencies. (n = 6-8 rats per light cycle) *** p < 0.0001

### Diurnal differences in baseline spontaneous firing frequency and firing patterns in CINs

Following voltammetry experiments, we wanted to target CINs directly via *ex vivo* electrophysiology. To maintain spontaneous firing of CINs, we utilized cell-attached recordings with aCSF as our internal solution (Fig. 2A). We also determined where in the NAc CINs were located after recordings and found a majority of CINs were in the NAc shell region (Fig. 2B). Next, diurnal variations were found in tonic CIN activity with significantly higher activity in the dark cycle in comparison to the light cycle (*p = 0.0003*) (Fig. 2C). Additionally, we were able to characterize CIN firing patterns into the following categories: regular, irregular, mixed mode, and bursting. These categories are based on previous research and take into consideration the coefficient of variation (CV) of the interspike interval (ISI) and pattern of spikes. There are no significant differences in the CV of ISI between the light and dark cycle (Fig. 2D). However, there are diurnal variations in the patterns of firing based on the spikes themselves (Fig. 3). During the light cycle, CINs exhibit more irregular firing patterns while CINs exhibit more regular firing in the dark cycle. Collectively, these data suggest that there are diurnal variations in CINs themselves that may contribute to the variations seen in NAc dynamics, like DA release.

**Figure 2.**
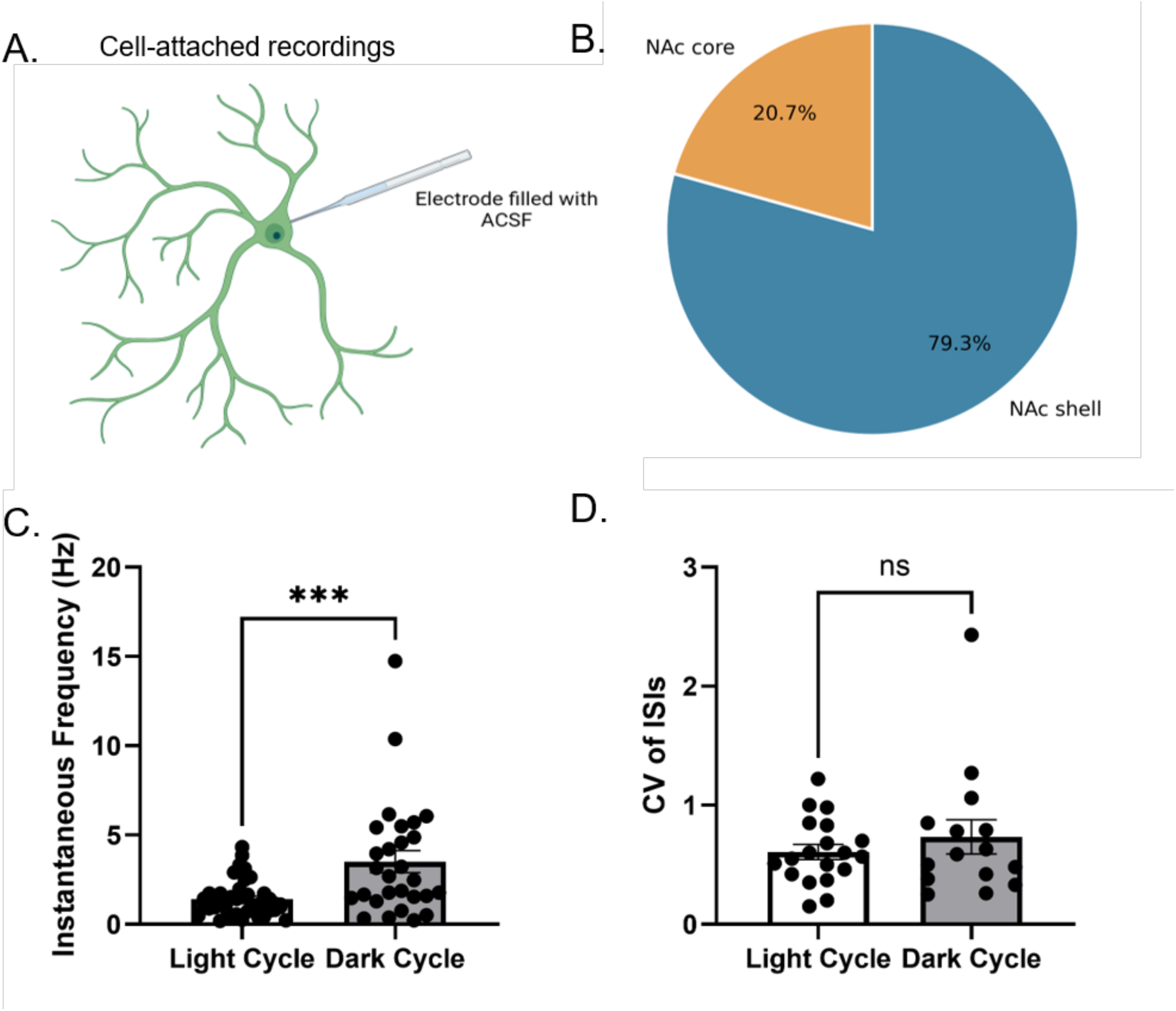
CINs in the NAc have higher firing frequencies during the dark cycle in comparison to the light cycle. (A) Cell-attach recordings were performed to measure the tonic activity of CINs without disrupting their spontaneous firing (B) All recordings took place in the NAc with a majority of CINs being in the NAc shell (C) Instantaneous firing frequency is higher in the dark cycle in comparison to the light (D) There are no significant differences between the light and dark cycle in the CV of ISIs (n = 4-6 mice with 1-2 cells per mouse per light cycle) *** p < 0.001

**Figure 3.**
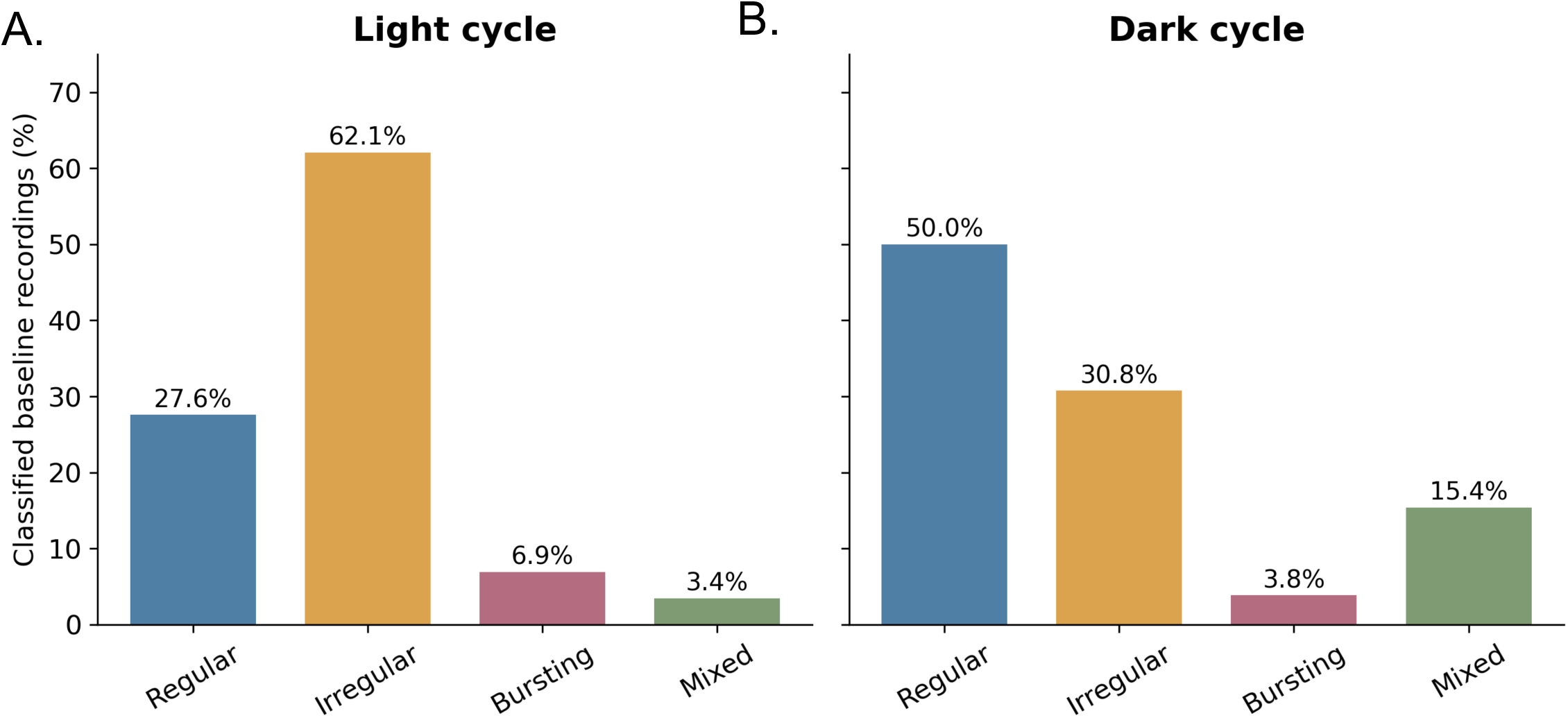
CIN firing patterns in the NAc vary between the light and dark cycle. (A) A majority of CIN firing patterns in the light cycle are classified as irregular (B) In contrast, most CIN firing patterns in the dark cycle are classified as regular (n = 4-6 mice with 1-2 cells per mouse per light cycle)

### Targeting nAChRs decreases spontaneous CIN firing frequency and firing patterns in the dark cycle

Similar to our voltammetry experiments, we wanted to determine the role of nAChRs in the diurnal variations in tonic CIN activity. We recorded CIN firing frequency after NIC bath application for 15 minutes after a 5-minute baseline. Two-way ANOVA revealed no significant main effects of light/dark cycle or drug treatment and no significant interaction between these factors (all p > 0.05). However, multiple comparisons revealed a significant reduction in CIN firing frequency following nicotine application during the dark cycle (*p = 0.0296*), but not during the light cycle (*p = 0.7172*) (Fig. 4A). Next, we washed on MEC for 15 minutes after a 5-minute baseline and found that there was no significant main effects of light/dark cycle or drug treatment and no significant interaction between these factors but there was a drug effect with significant a decrease in both the light (*p = 0.0466*) and dark (*p = 0.0217*) cycle (Fig. 4B). Lastly, we utilized DHβE without CON to target specific nAChRs that contribute to DA release and found that there was an effect of light cycle with activity being higher in the dark *(p = 0.0016)* the dark cycle in comparison to the light cycle and that DHβE significantly decreased CIN firing in the dark cycle *(p = 0.0485)* (Fig. 4C). We found overall that targeting nAChRs mostly had effects in the dark cycle while only MEC decreased firing in the light cycle. Interestingly, we only find changes in the firing patterns in the dark cycle after NIC (Fig. 5D) and MEC (Fig. 5E) wash on. Overall, these data demonstrate that targeting nAChRs decreases CIN firing primarily in the dark cycle while also changing firing patterns.

**Figure 4.**
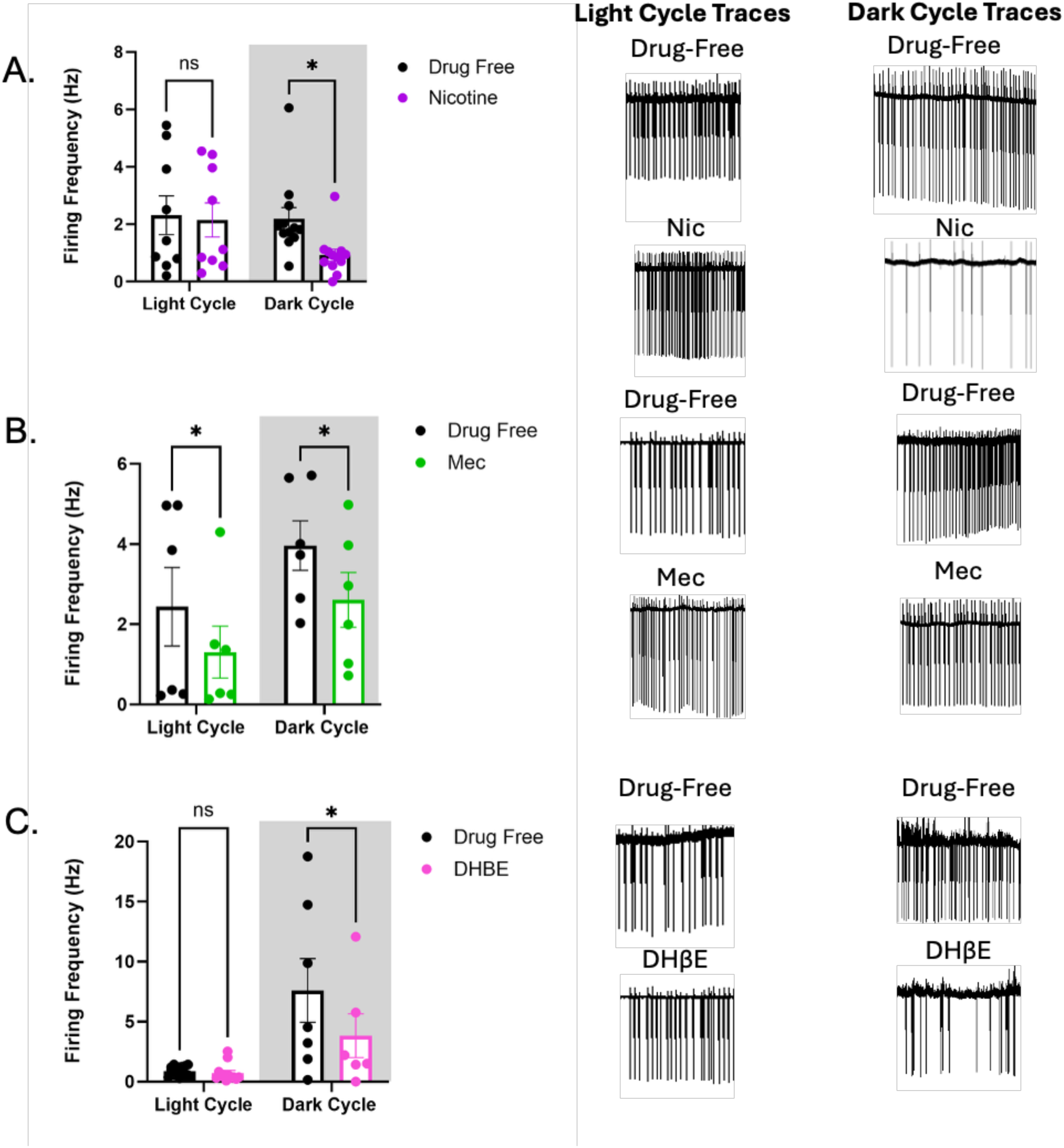
Diurnal variations in the effects of nAChRs on CIN firing. (A) Bath application of NIC, a nAChR agonist, decreases CIN firing frequency in the dark cycle in comparison to the light cycle (B) Interestingly, MEC, a general nAChR antagonist, decreases firing in both the light and dark cycle CIN firing (C) DHBE, an antagonist targeting beta 2 containing nAChR decreased CIN firing in the dark cycle in comparison to the light cycle (n = 4-6 mice with 1-2 cells per mouse per light cycle) * p < 0.05

**Figure 5.**
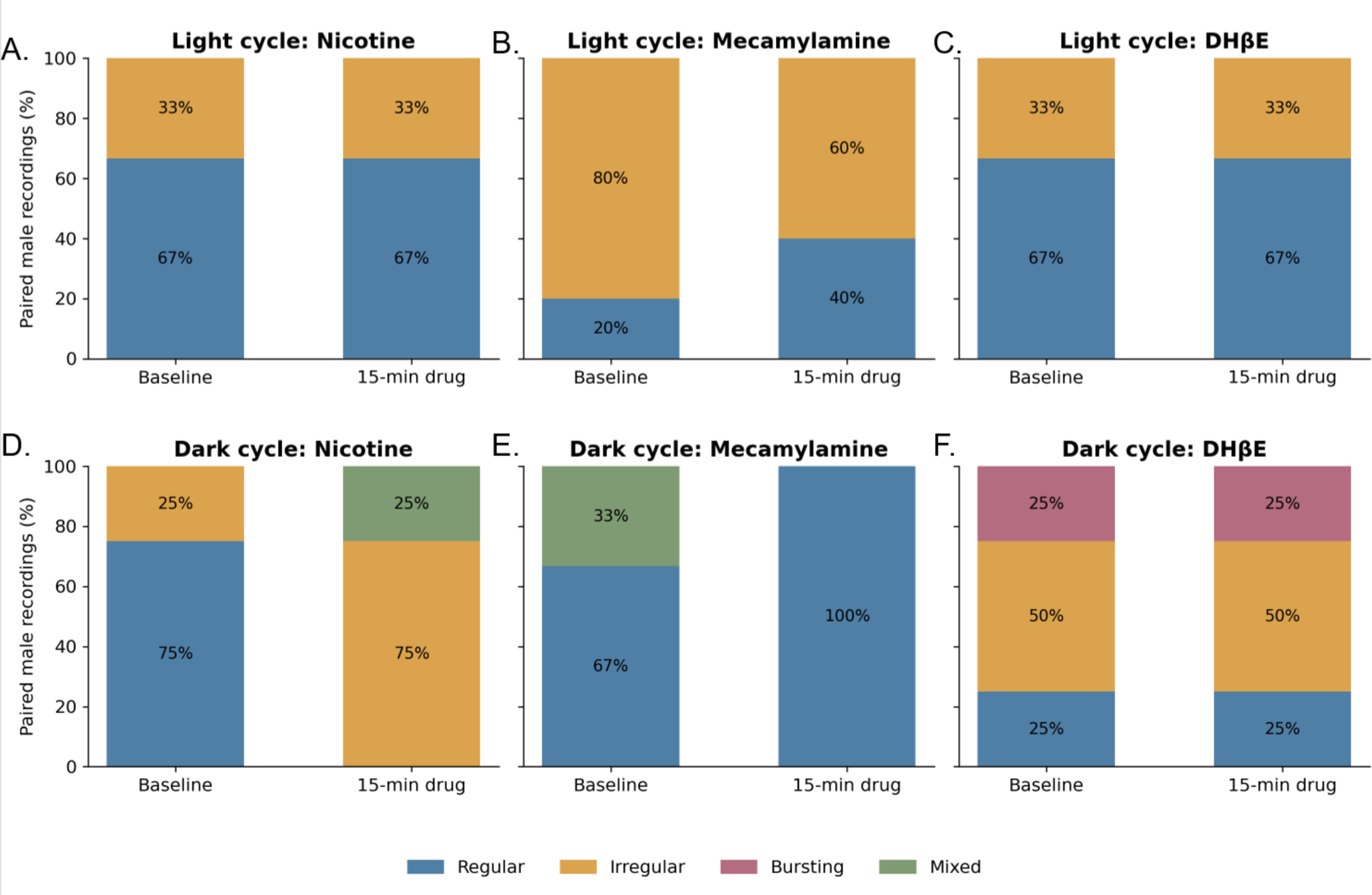
Broad targeting of nAChRs changes firing patterns of CINs exclusively in the dark cycle. (A) Bath application of drugs targeting nAChRs had little to know effects on firing patterns in the light cycle (B) Bath application of NIC and MEC changed the firing patterns of CINs while DHBE had little to know effects on firing patterns, which suggest that targeting nAChR regardless of subunit may have more of an effect on firing patterns in the dark cycle (n = 4-6 mice with 1-2 cells per mouse per light cycle)

## Discussion

Here, we identified and determined diurnal variations in CIN regulation within the NAc by examining the effects of nAChR-targeting drugs on stimulated DA release across varying stimulation frequencies and spontaneous CIN activity. Previous studies demonstrated diurnal variations in stimulated DA release and its modulation by CIN-derived ACh using *ex vivo* optogenetics⁹. In the present study, the effects of nAChR-targeting drugs on stimulated DA release, normalized to baseline, did not differ between the light and dark phases. In contrast, these drugs altered the spontaneous firing frequency of CIN activity primarily during the dark phase. Together, these findings suggest that time of day may influence the sensitivity of CIN activity to nAChR-targeting drugs while other mechanisms may play more of role in the diurnal variations in stimulated DA release.

Diurnal variations occur in the neurobiology underlying reward-related behaviors, including neuronal activity and DA signaling within the NAc. For example, NAc MSNs exhibit greater excitability during the dark phase than the light phase¹⁵, while extracellular DA tone, stimulated DA release, and DA uptake vary across the light/dark cycle⁹˒¹⁴. Previous work also demonstrated diurnal differences in the modulation of DA release by CIN-derived ACh⁹. In the present study, pharmacological modulation of nAChRs did not produce significant phase-dependent differences in stimulated DA release when expressed as a percentage of baseline. However, MEC produced a larger reduction in stimulated DA release during the light cycle, with minimal effects observed during the dark cycle. This finding is consistent with our electrophysiological data demonstrating decreased CIN firing following MEC application during the light cycle. Together, these findings suggest that nicotinic signaling contributes to the regulation of both cholinergic and dopaminergic activity, with potential differences across the light/dark cycle.

Interestingly, NIC and MEC produced modest increases in stimulated DA release at higher stimulation frequencies during the dark cycle, whereas reductions were observed across frequencies during the light cycle. In contrast, DHβE, which preferentially antagonizes β2-containing nAChRs, had minimal effects during either phase. Previous work demonstrated that NIC increased the phasic/tonic ratio of stimulated DA release to a greater extent during the dark phase⁹. These findings suggest that the effects of nicotinic modulation on DA release may depend on stimulation frequency, nicotine concentration, and time-of-day. Additionally, other local mechanisms, including D2 receptor-mediated regulation of CIN activity, may contribute to diurnal variations in NAc cholinergic signaling. Further studies are needed to determine how these receptor systems interact to regulate CIN activity and DA release across the light/dark cycle.

When examining spontaneous CIN firing in the NAc, we found higher firing frequency during the dark phase in comparison to the light phase. This finding suggests that CIN signaling may vary across the day, with increased CIN firing potentially supporting greater ACh output during the rodent’s active phase. Although ACh release was not measured directly, this interpretation is consistent with previous findings of higher stimulated DA release and phasic/tonic release ratios midway through the dark phase⁹˒¹⁴. Together, these data suggest coordinated diurnal regulation of CIN activity and DA signaling within the NAc, which may contribute to time-of-day differences in reward-related behaviors.

In addition to firing frequencies, we also found diurnal variations in the firing patterns. A majority of CINs in the light cycle exhibited irregular firing patterns while CINs in the dark cycle mostly exhibit regular firing patterns. Various firing patterns are a unique characteristic of CINs as they are spontaneously active even in slices^9,23^. Additionally, the firing patterns of CINs play a role in reward-related behaviors. For example, enhancing the bursting of CINs increases extinction while disrupting the pauses in firing disrupts goal-directed behaviors ^24^. Thus, diurnal variations in CIN firing patterns may relate to the diurnal variations seen in reward-related behavior.

CINs that fire more regularly tend to fire faster, which may explain why CINs in the dark cycle have both higher frequencies and regular firing in comparison to CINs in the light cycle^25^. During the light cycle, most CINs exhibited irregular firing patterns with some bursting. Limited studies have examined the diurnal variations in goal-directed and reward-directed behaviors, but data show that goal-directed behaviors are predominant in the light cycle^10^. Additionally, when examining just ACh modulation of DA release, it was higher in the light cycle, which further supports that the diurnal variations in local NAc dynamics may drive different behaviors.

Next, we found a decrease in CIN firing following NIC, MEC, and DHβE application during the dark cycle while only MEC impacting CIN firing in the light cycle. These findings suggest that endogenous nicotinic signaling contributes to maintaining CIN activity during the rodent’s active phase. Although nicotine activates nAChRs, sustained exposure can promote receptor desensitization, providing a possible explanation for why NIC and nicotinic antagonists produced similar reductions in firing. Consistent with this, previous work demonstrated that bath-applied nicotine reduced spontaneous CIN firing in the dorsal striatum^26^. However, the mechanisms underlying these effects in the NAc remain unknown, as bath application can influence receptors on CINs and on other elements of the local circuit.

Additionally, NIC and MEC alter CIN firing patterns during the dark cycle, whereas DHβE reduces firing frequency without altering firing patterns. DHβE preferentially antagonizes β2-containing nAChRs, particularly α4β2 receptors, which are expressed on dopaminergic terminals and other neuronal populations within the striatum. In contrast, nicotine and mecamylamine act on a broader range of nAChR subtypes, potentially influencing additional components of the local circuitry that contribute to CIN firing patterns. These findings suggest that distinct nAChR populations or downstream circuit mechanisms may differentially regulate CIN firing frequency and firing patterns. Further investigation is needed to determine how these mechanisms contribute to diurnal variations in NAc CIN signaling.

These data are the first to exhibit diurnal variations in CIN firing patterns and the impact of nAChRs on diurnal variations in CIN firing frequency and patterns. To measure tonic activity of CINs, we utilized cell-attached electrophysiology to avoid disrupting the spontaneous firing patterns. Cell-attached recordings are commonly used in spontaneously active cells, but do not give us specifics about the cell, like resting membrane potential^27^. Future studies will include whole-cell recordings as well along with other specific techniques, like *ex vivo* optogenetics, to potentially parse out mechanisms driving diurnal variations.

Although these initial studies expand our understanding of diurnal variations in the CIN system in the NAc, there remain limitations. Our current studies utilize both rat and mouse models to determine different aspects of the NAc CIN system. Rats were used for ex vivo voltammetry experiments to examine stimulated DA release and its modulation by nAChRs within the NAc. In contrast, mice were used for electrophysiological recordings to take advantage of a transgenic line that allowed for the direct identification of CINs. Since CINs make up about 1–2% of neurons in the NAc, identifying these cells based on morphology alone would be challenging and could potentially lead to misidentification. Thus, the use of both species allowed us to examine distinct but complementary components of the NAc CIN system. Future studies will aim to utilize transgenic mouse models across experimental approaches to reduce potential species-related variability and allow for more targeted approaches in the NAc circuitry. Moreover, we only utilized one dose of each drug based on literature and our previous work. Future studies will investigate additional doses and determine if there is a dose dependent effect on the diurnal variations in CINs. Lastly, we focused on male subjects and may be missing potential sex differences. Limited studies have found that hormones may play a key role in CIN firing^28^. Future studies will focus on the potential sex differences in the CIN system that may also have varying diurnal differences.

Overall, our findings enhance understanding of diurnal regulation within the NAc CIN system and suggest that time-of-day is an important consideration when examining CIN contributions to reward-related signaling. Differences in spontaneous CIN firing and nAChR mediation may contribute to the temporal regulation of local DA dynamics. Determining these baseline variations provides a foundation for determining how CIN function is altered by repeated drug exposure and contributes to SUDs. Specifically, this may be relevant to nicotine, which activates nAChRs involved in endogenous CIN signaling, and NAc CINs have been implicated directly in nicotine seeking^29^. Human studies also demonstrate time-of-day variation in smoking and craving. For example, ecological momentary assessment identified elevated smoking during the first two hours after waking and a smaller evening peak. These data support that there are diurnal differences in NAc CIN function that may influence responses to nicotine and nicotine-associated cues. Our future studies aim to incorporate preclinical models of SUDs in combination with CIN manipulation to determine how the diurnal variations in CINs contribute to behaviors, like drug-taking.

## Acknowledgments

The author(s) declare that financial support was received for the research, authorship, and/or publication of this article. This work was funded by the National Institutes of Health: DA039865 (PI: Colleen McClung) and DA060613 (PI: Taylor Stowe).

## Conflicts of interest

The author(s) have no conflicts of interest to disclose.

## Notes

### Competing Interest Statement

The authors have declared no competing interest.

